# Flex ddG: Rosetta ensemble-based estimation of changes in protein-protein binding affinity upon mutation

**DOI:** 10.1101/221689

**Authors:** Kyle A. Barlow, Shane O Conchúir, Samuel Thompson, Pooja Suresh, James E. Lucas, Markus Heinonen, Tanja Kortemme

## Abstract

Computationally modeling changes in binding free energies upon mutation (interface ΔΔ*G*) allows large-scale prediction and perturbation of protein-protein interactions. Additionally, methods that consider and sample relevant conformational plasticity should be able to achieve higher prediction accuracy over methods that do not. To test this hypothesis, we developed a method within the Rosetta macromolecular modeling suite (flex ddG) that samples conformational diversity using “backrub” to generate an ensemble of models, then applying torsion minimization, side chain repacking and averaging across this ensemble to estimate interface ΔΔ*G* values. We tested our method on a curated benchmark set of 1240 mutants, and found the method outperformed existing methods that sampled conformational space to a lesser degree. We observed considerable improvements with flex ddG over existing methods on the subset of small side chain to large side chain mutations, as well as for multiple simultaneous non-alanine mutations, stabilizing mutations, and mutations in antibody-antigen interfaces. Finally, we applied a generalized additive model (GAM) approach to the Rosetta energy function; the resulting non-linear reweighting model improved agreement with experimentally determined interface DDG values, but also highlights the necessity of future energy function improvements.

## Introduction

Protein-protein interactions underlie essentially all biological processes, including signal transduction and antibody-antigen recognition. Many protein-protein interfaces are sensitive to mutations that can alter interaction affinity and specificity. In fact, mutations at protein-protein interfaces have been reported to be overrepresented within disease-causing mutations,^1^ highlighting the central importance of these interactions to biology and human health. A sufficiently accurate computational method capable of predicting mutations that strengthen or weaken known protein-protein interactions would hence serve as a useful tool to dissect the role of specific protein-protein interactions in important biological processes. Coupled with state-of-the-art methods for protein engineering and design, such a method would also enhance our ability to create new and selective interactions, enabling the development of improved protein therapeutics, protein-based sensors, and protein materials.

Several prior methods have been developed to predict changes in protein-protein binding affinity upon mutation using different approaches to estimating energetic effects (scoring) and modeling structural changes (sampling). Common approaches include weighted energy functions that seek to describe physical interactions underlying protein-protein interactions,^2,3^ statistical and contact potentials,^4–7^ a combination of these approaches,^8,9^ graph-based representations,^10^ methods that sample backbone structure space locally around mutations,^11^ and machine learning approaches,^12^

We set out to develop and assess methods for estimating experimentally determined changes in binding free energy after mutation (interface ΔΔ*G*) within the Rosetta macro-molecular modeling suite, Rosetta is freely available for academic use, and allows combination of interface ΔΔ*G* predictions with Rosetta’s powerful protein design capabilities, which have proven successful in a variety of applications,^13,14^ Prior projects have applied Rosetta predictions to dissect determinants of binding specificity and promiscuity,^15,16^ enhance protein-protein binding affinities,^17,18^ and to design modified^19^ and new interactions,^20–22^ but no prior benchmarking effort has quantitatively assessed the performance of predicting changes in binding free energy in Rosetta on a large, diverse benchmark dataset, in part because such datasets have only become available more recently. The current state-of-the-art Rosetta ΔΔ*G* method, ddg_monomer,^23^ has proven effective at predicting changes in stability of monomeric proteins after mutation, but had not yet been tested at predicting change of binding free energies in protein-protein complexes. Prior “computational alanine scanning” ΔΔ*G* methods were benchmarked on mutations in protein-protein interfaces, focusing on mutations to alanine,^24–26^ The original Rosetta alanine scanning method^24^ did not sample backbone degrees of freedom, which is a first-order approximation for mutations to alanine (that are not expected to cause large backbone perturbations^27^), but less likely to be predictive for mutations to larger side chains which might require some degree of backbone rearrangement to accommodate the change. Inclusion of recent Rosetta energy function and sampling method developments, including methods that attempt to more aggressively sample conformational space, has not resulted in significant improvement to the alanine scanning method.^26^

We sought to create a method that would take into account aspects of the conformational plasticity of proteins by representing structures as an ensemble of individual full-atom models to explore biologically relevant and accessible portions of conformational space near the crystallographically determined input structures. Ensemble representations have previously been shown to be effective at predicting changes in protein stabilities after mutation and at predicting the effects of mutation on protein-protein binding affinities,^28^ as well as at improving Δ*G_binding_* calculations between kinases and their inhibitors.^29^

We chose to sample conformational plasticity using the “backrub” protocol implemented in Rosetta.^30^ The backrub method samples local side chain and backbone conformational changes, similar to those suggested to underlie observed conformational heterogeneity in high-resolution crystal structures,^31^ and to accommodate evolved and designed mutations.^32^ Backrub ensembles have been demonstrated to recapitulate properties of proteins that have been experimentally determined, such as side chain NMR order parameters,^33^ tolerated sequence profiles at protein-protein^34^ and protein-peptide interfaces,^35,36^ and conformational variability between protein homologs.^37^ Backrub has also proved effective in design applications, such as the redesign of protein-protein interfaces^19^ and recapitulation of mutations that alter ligand-binding specificity.^38^ When compared to ensembles generated via molecular dynamics simulations or the “PertMin” method,^39^ backrub ensembles were shown to be the only ensembles capable of generating higher diversity (as measured by RMSD) between output models than from output models to the original input crystal structure. This observation suggests that backrub could be uniquely suited to produce diverse ensembles that effectively explore the local conformational space around an input structure.^39^ Taken together, we hypothesized that these previously demonstrated properties of backrub ensembles would also make them an effective representation of near-native conformational states for use in predicting interface ΔΔ*G* values.

## Methods

### Benchmark datasets

Developing and assessing the accuracy of a new method to predict changes in binding free energy after mutation requires a large and diverse benchmark set covering single mutations to all amino acid types, multiple mutations, and mutations across a variety of protein-protein interfaces. To facilitate comparisons to other methods and to avoid biases specific to our approach, we chose to use an existing benchmark dataset created by Dourado and Flores^11^ during the development of their ZEMu (Zone Equilibration of Mutants) method. The ZEMu dataset was curated from the larger SKEMPI database^40^ by avoiding a bias towards complexes in which a single position is repeatedly mutated, experimental data that are not peer-reviewed, redundancy (duplicate experimental values), mutations outside of interfaces, mutations involved in crystal contacts, and experimental ΔΔ*G* values for which wild-type and mutant conditions (such as pH) varied. Confidence in the “known” experimental ΔΔ*G* values is important, as it has been pointed out that the experimental methodology used can have a strong effect on the performance of predictors of changes in binding free energy.^41^ The ZEMu dataset was also curated to include a range of both stabilizing and destabilizing mutants, small side chain to large side chain mutations, single and multiple mutations, and a diversity of complexes. Small-to-large mutations are defined as those dataset cases where all mutation(s) are at positions where the residue side chain increases in van der Waals volume post-mutation.^42^

After a review of the literature from which the known experimental ΔΔ*G* values originated, we removed one data point from the 1254 point ZEMu set that we could not match to the originally reported affinity value. We also removed 5 mutations we determined to be duplicates, along with 8 mutations that were reverse mutations of other data points, leaving us with a test set of 1240 mutations (Table 1). We used SAbDab^43^ to define complexes that contained at least one antibody binding partner. Our version of the ZEMu dataset is available in the Supporting Information as Dataset S1. All ΔΔ*G* predictions described in the paper are available in the Supporting Information.

**Table 1:**
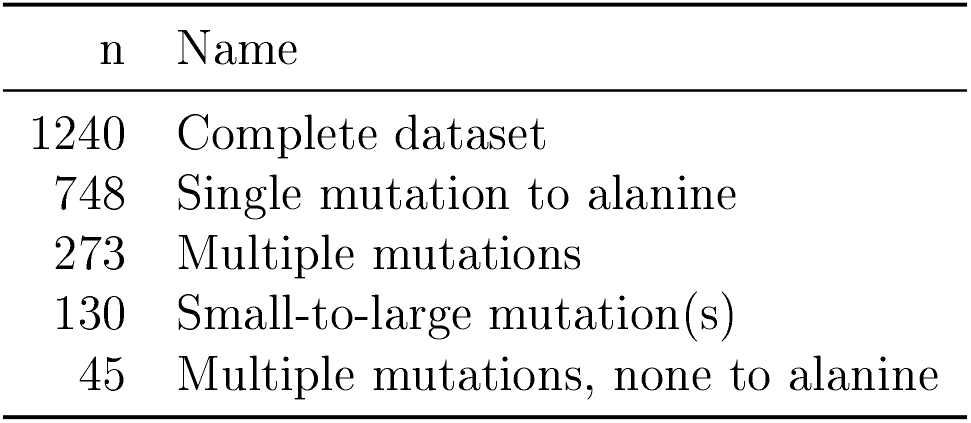
ZEMu dataset composition

### Rosetta implementation and prediction protocol

Our protocol, called “flex ddG”, is implemented within the RosettaScripts interface to the Rosetta macromolecular modeling software suite,^44^ which makes the protocol easily adaptable to future improvements and energy function development. The method can be run using a Rosetta Scripts XML that is available in the Supporting Information as Listing 1. Version numbers of tested software are available in Table S1.

Flex ddG method steps are outlined in Fig. 1. **Step 1:** The protocol begins with an initial minimization (on backbone *ϕ*/*ψ* and side chain *χ* torsional degrees of freedom, using the “lbfgs_armijo_nonmonotone” minimization algorithm option) of the input crystal structure of the wild-type protein complex. This (and later) minimizations are performed with constraints that harmonically restrain pairwise atom distance to their values in the input crystal structure. Minimization is run until convergence (absolute score change upon minimization of less than one REU (Rosetta Energy Unit)). **Step 2:** Starting from the minimized input structure, the backrub method in Rosetta^30^ is used to create an ensemble of models. In brief, each backrub move is undertaken on a randomly chosen protein segment consisting of three to twelve adjacent residues in the neighborhood of any mutated position, The mutation neighborhood is defined by finding all residues with a C-*β* atom (C-*α* for glycines) within 8 Å of any mutant position, then adding this residue and its adjacent N and C-terminal residues to the list of neighborhood residues. All atoms in the backrub segment are rotated locally about an axis defined as the vector between the endpoint C-*α* atoms. Backrub is run at a temperature of 1.2 kT, for up to 50,000 backrub Monte Carlo trials/steps (Table S2 shows that using a kT of 1.6 gives similar results to a kT of 1.2). Up to 50 output models are generated. **Step 3A:** For each of the 50 models in the ensemble output by backrub, the Rosetta “packer” is used to optimize side chain conformations for the wild-type sequence using discrete rotameric conformations^45^ and simulated annealing. The packer is run with the multi-cool annealer option,^46^ which is set to keep a history of the 6 best rotameric states visited during annealing. **Step 3B:** Independently and in parallel to step 3A, side chain conformations for the mutant sequence are optimized on all 50 models, introducing the mutation(s). **Step 4A:** Each of the 50 wild-type models is minimized, again adding pairwise atom-atom constraints to the input structure. Minimization is run with the same parameters as in step 1; the coordinate constraints used in this step are taken from the coordinates of the Step 3A model. **Step 4B:** As Step 4A, but for each of the 50 mutant models. **Step 5A:** Each of the 50 minimized wild-type models are scored in complex, and the complex partners are scored individually. The scores of the split, unbound complex partners are obtained simply by moving the complex halves away from each other. No further minimization or side chain optimization is performed on the unbound partners before scoring. **Step 5B:** In the same fashion as Step 5A, each of the 50 minimized mutant models are scored in complex, and the complex partners are scored individually. **Step 6:** The interface ΔΔ*G* score is produced via Eq. 1:

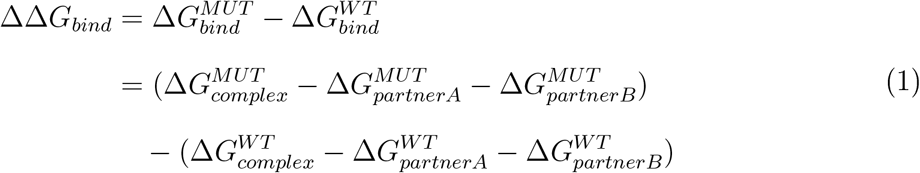

**Figure 1:**
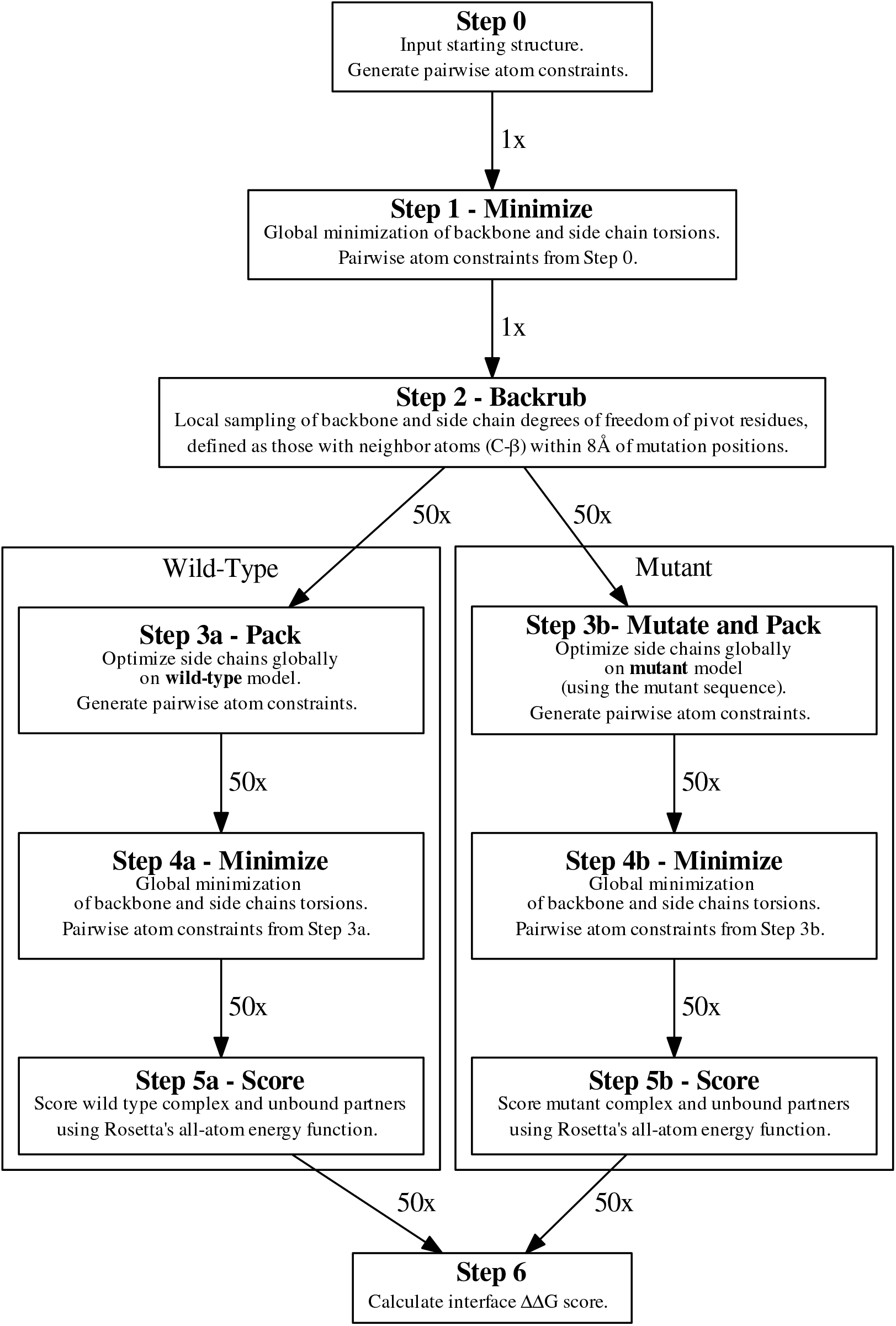
Schematic of the flex ddG protocol method.

We evaluate performance of the protocol by comparing predicted ΔΔ*G* scores to known experimental values, using Pearson’s correlation (R), Fraction Correct (FC), and Mean Absolute Error (MAE). Fraction Correct is defined as the number of cases in the dataset categorized correctly as stabilizing, neutral, or destabilizing, divided by the total number of cases in the dataset. Stabilizing mutations are defined as those with a ΔΔ*G* <= −1.0 kcal/mol, neutral as those with −1.0 kcal/mol < ΔΔ*G* < 1.0 kcal/mol, and destabilizing as those with ΔΔ*G* >= 1.0 kcal/mol.

MAE (Mean Absolute Error) is defined in Eq. 2 as:

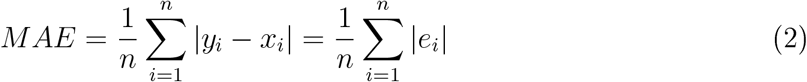

where *y_i_* are the predicted ΔΔ*G* values, *x_i_* are the known, experimentally determined values, and *e_i_* is the prediction error.

### Rosetta energy function

We utilized Rosetta’s Talaris^45,47,48^ all-atom energy function for the modeling steps. As we do not modify our models of the unbound state, several terms of the Rosetta energy function will cancel out in the final ΔΔ*G* scoring because the Δ*G* of folding score of the unbound partners is subtracted from the total score of the complex (Eq. (1)). After subtraction, seven score terms remain, and combined, become the final interface ΔΔ*G* score, dominated by solvation (fa_sol using an implicit solvation model^49^), hydrogen bonding and electrostatics^47,48,50^ (hbond_sc: side chain-side chain hydrogen bonds; hbond_bb_sc: hydrogen bonds between backbone atoms and side chain atoms; hbond_lr_bb: long-range hydrogen bond interactions between backbone atoms; fa_elec: Coulomb electrostatics), and Lennard-Jones atomic packing interactions (fa_rep and fa_atr: repulsive and attractive components of the Lennard-Jones potential).

### Score analysis

To investigate potential sources of prediction error on an individual score term basis, we used a generalized additive model^51^ approach to fit Rosetta’s predicted ΔΔ*G* values to experimentally known values. First, we apply an unbiased logistic scaling to individual score terms,

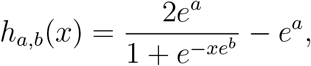

where *a* is the scaling range of the score, and *b* is the steepness of the sigmoid scaling. Both parameters are transformed through an exponential to ensure non-negativity. The scaling function *h* does not introduce bias, that is, *h_θ_*(0) = 0 for any *θ*, The scoring model results in a generalized additive model (GAM) over the *M* score terms,

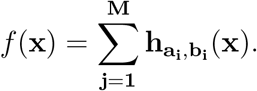

The parameters 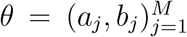 for the score terms were simultaneously sampled using a random walk Metropolis-Hastings MCMC algorithm (the `mhsample` function in Matlab) assuming a Gaussian likelihood as the target distribution

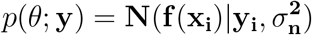

with a noise variance set to 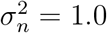, and where 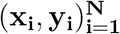 are the empirical observations *y_i_* that correspond to the protein score terms **x_i_**, respectively. We sample for 1000 samples with a burn-in set to 1000 samples and a thinning parameter of 20. The proposal distribution was selected to be a symmetric uniform distribution such that [*a*^(*s*+1)^, *b*^(*s*+1)^] ~ *U*(*a*^(*s*)^ ± 2, *b*^(*s*)^ ± 2). The resulting MCMC sample represents all logistics score scalings that reproduce the empirical measurements assuming an error model with noise variance 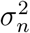.

## Results and discussion

The overall performance of the protocol is summarized in Table 2. We compare 4 prediction methods: (a) our flex ddG backrub ensemble method, (b) the prior state-of-the-art Rosetta methodology, ddg_monomer,^23^ (c) a control version of our flex ddG protocol which omits the backrub ensemble generation step, leaving only the minimization and packing steps, and (d) published data from the ZEMu (zone equilibration of mutants) method.^11^ Data split by input protein-protein complex are shown in Table S3.

**Table 2:** Summary of prediction performance. Flex ddG predictions used 50 models and 35000 backrub steps. ddG monomer predictions used the default of averaging the ΔΔ*G* scores of the three lowest scoring output models, as implemented in the original method.^23^ N = number of cases in the dataset or subset. R = Pearson’s R. MAE = Mean Absolute Error. FC = Fraction Correct. Best performance for each metric and dataset is shown in bold.

| Mutation Category | Prediction Method | N | R | MAE | FC |
| --- | --- | --- | --- | --- | --- |
| Complete dataset | flex ddG | 1240 | **0.63** | **0.96** | **0.76** |
|  | ddG monomer |  | 0.51 | 1.57 | 0.64 |
|  | no backrub control |  | 0.56 | 1.12 | 0.73 |
|  | ZEMu paper |  | 0.61 | 1.08 | 0.71 |
| Small-to-large mutation(s) | flex ddG | 130 | **0.65** | **0.78** | **0.71** |
|  | ddG monomer |  | 0.31 | 1.55 | 0.55 |
|  | no backrub control |  | 0.41 | 1.11 | 0.62 |
|  | ZEMu paper |  | 0.48 | 1.16 | 0.65 |
| Mutation(s) to alanine | flex ddG | 939 | **0.62** | **0.96** | **0.78** |
|  | ddG monomer |  | 0.50 | 1.55 | 0.66 |
|  | no backrub control |  | 0.58 | 1.06 | 0.75 |
|  | ZEMu paper |  | **0.62** | 1.03 | 0.73 |
| Single mutation to alanine | flex ddG | 748 | **0.51** | **0.75** | **0.76** |
|  | ddG monomer |  | 0.36 | 1.31 | 0.62 |
|  | no backrub control |  | 0.44 | 0.90 | 0.74 |
|  | ZEMu paper |  | 0.45 | 0.86 | 0.71 |
| Multiple mutations | flex ddG | 273 | 0.62 | **1.62** | **0.78** |
|  | ddG monomer |  | 0.50 | 2.44 | 0.70 |
|  | no backrub control |  | 0.58 | 1.73 | 0.73 |
|  | ZEMu paper |  | **0.64** | 1.63 | 0.75 |
| Multiple mutations, all to alanine | flex ddG | 191 | 0.47 | 1.77 | **0.84** |
|  | ddG monomer |  | 0.34 | 2.49 | 0.80 |
|  | no backrub control |  | 0.50 | **1.69** | 0.81 |
|  | ZEMu paper |  | **0.55** | 1.72 | 0.79 |
| Multiple mutations, none to alanine | flex ddG | 45 | **0.63** | **1.38** | **0.60** |
|  | ddG monomer |  | 0.40 | 2.54 | 0.38 |
|  | no backrub control |  | 0.44 | 1.66 | 0.58 |
|  | ZEMu paper |  | 0.53 | 1.59 | **0.60** |
| Antibodies | flex ddG | 355 | **0.61** | **0.93** | **0.74** |
|  | ddG monomer |  | 0.50 | 1.35 | 0.69 |
|  | no backrub control |  | 0.49 | 1.06 | 0.72 |
|  | ZEMu paper |  | 0.54 | 1.06 | 0.67 |

The new flex ddG method outperforms the comparison methods on the complete dataset in each of the correlation, MAE, and fraction correct metrics (Table 2). In particular, we see a large increase in performance relative to the other methods on the small-to-large subset of mutations. This is in accordance with our expectations that backrub ensembles should be able to sample small backbone conformational adjustments required to accommodate changes in amino acid residue size. Notably, application of backrub ensembles performs better than other methods that include backbone minimization steps only, including the current state-of-the-art Rosetta ddg_monomer method. On the small-to-large mutations subset, the ddg_monomer method achieves a Pearson correlation of only 0.31 compared to 0.65 with flex ddg.

Performance of the flex ddG method on the subset of single mutations to alanine is also competitive or outperforms the alternative methods. As we do not expect single mutations to alanine to require intensive backbone sampling, our method’s effectiveness on this subset shows that the method is fairly robust to the mutation type. As we chose to perform backrub sampling prior to introducing mutations, these results could suggest that flex ddG is effective by sampling underlying, relevant plasticity of the input crystal structure instead of distorting the local structure around a mutation to resolve a clash or poor interaction with a mutant side chain.

While the flex ddG method shows improved performance on the subset of multiple mutations as compared to the control and ddg_monomer methods, flex ddg did not match the performance of the ZEMu method on this subset. This result could indicate that further refinement of the backrub parameters is required when simultaneously sampling conformational space around the sites of multiple mutations. However, and remarkably, flex ddG outperforms ZEMu on the subset of cases with multiple mutations where none of the mutations are to alanine (Table 2). Finally, the flex ddg method also shows considerable improvements over other methods on the subset of antibody-antigen complexes (Table 2).

Fig. 2 illustrates the performance for the flex ddG and control methods on the complete dataset and small-to-large subsets using scatterplots comparing experimentally determined and computationally estimated changes in binding free energies for each of the cases in the datasets. In particular, a notable improvement with flex ddG over the control can be seen for the 13 small-to-large mutations that were experimentally determined to stabilize the protein-protein interface significantly (ΔΔ*G* <= −1.0 keal/mol). For this set, the control method misclassifies most stabilizing mutations to have minimal effect or to be destabilizing (9 mutations with predicted Rosetta ΔΔ*G* scores > 0) (Fig. 2d), whereas flex ddG identifies a sizable number (12 of 13 mutations) to have predicted Rosetta ΔΔ*G* < 0 (Fig. 2c), even though only one of these mutations is predicted to be strongly stabilizing (predicted ΔΔ*G* score < 1). The capability to predict stabilizing mutations is especially important for challenging design applications to modulate binding affinity and selectivity, as well as creating entirely new high-affinity protein-protein interactions.

**Figure 2:**
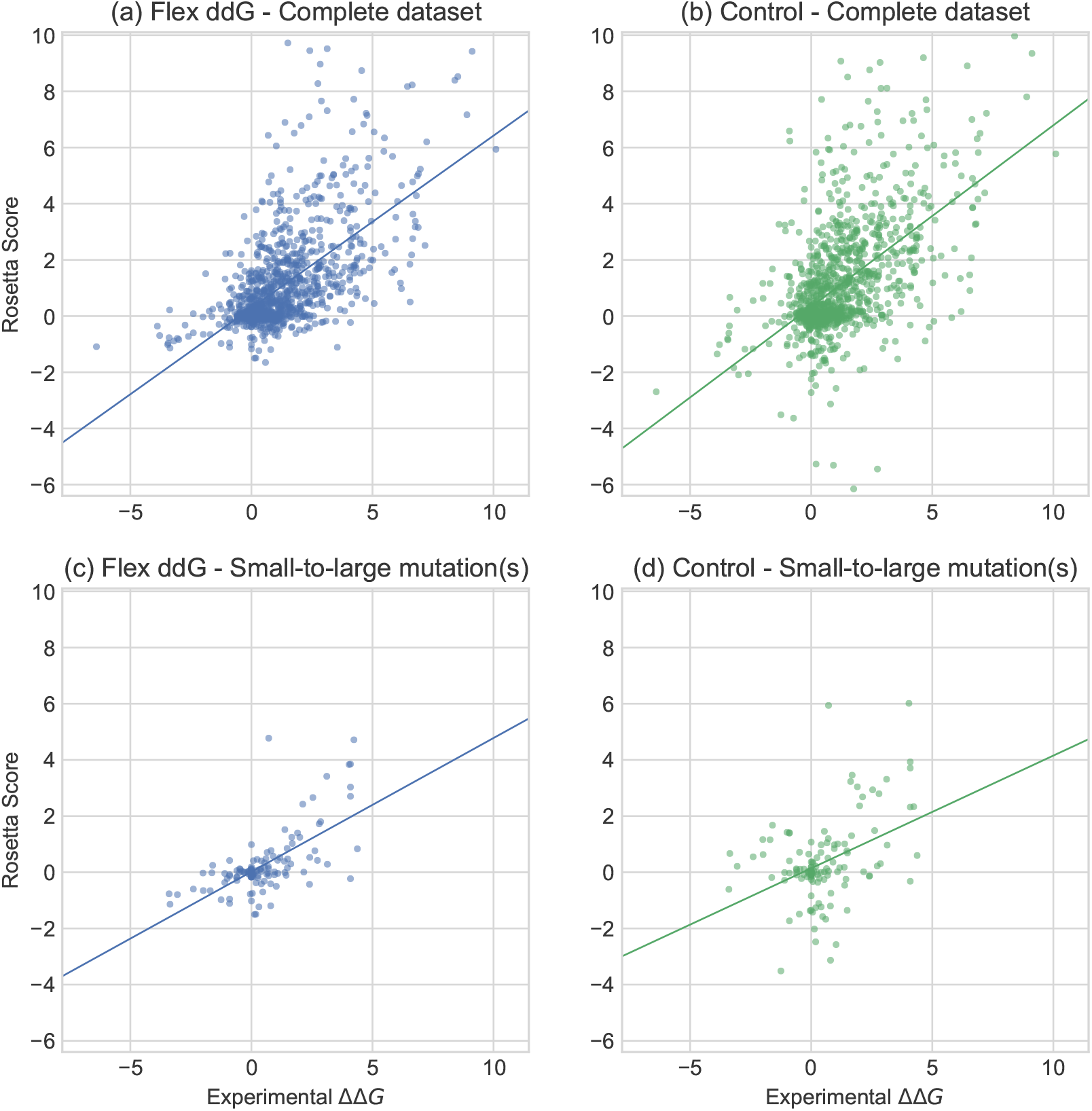
Experimentally determined ΔΔ*G* values (x-axis) versus Rosetta predictions. Rosetta scores are in Rosetta Energy Units (REU) using the Rosetta Talaris energy function,^45,47,48^ (a) flex ddG method (35000 backrub steps); Complete dataset (n=1240). (b) no backrub control; Complete dataset (n=1240). (c) flex ddG method (35000 backrub steps); Small-to-large mutation(s) (n=130). (d) no backrub control; Small-to-large mutation(s) (n=130).

In the following sections, we assess how different flex ddG implementations would affect prediction performance, focusing separately on sampling and scoring.

### Effect of ensemble size

While the results presented above used an ensemble size of 50 members, we next investigated what the ideal ensemble size would be to maximize the predictive ability of our method. For example, prior methods used ensemble sizes ranging from ten^3^ to thousands.^28^ As the computational time required to run flex ddG increases linearly with ensemble size, determining an optimal size is practically relevant. We therefore evaluated the performance of flex ddG as we average across an increasing number of models (from 1 to 50, Fig. 3). The models are first sorted by the score of the corresponding repacked and minimized wild type model, such that producing a ΔΔ*G* with 1 model will only use the lowest (best) scoring model, 2 models will use the 2 lowest scoring models, and so forth. Fig. 3(a) shows the performance on the complete dataset. As more models with increasing wild type complex score are averaged, correlation with known experimental values increases. Conversely, performance for the no backrub control method stays approximately constant as more models are averaged. This result indicates that sampling with backrub adds information that improves ΔΔ*G* calculation even though the additional averaged models have higher scores (average ensemble total score is shown in Fig. S1). These higher scoring models would be excluded in methods such as the Rosetta ddg_monomer protocol, which typically use only the lowest scoring wild-type and mutant models.

**Figure 3:**
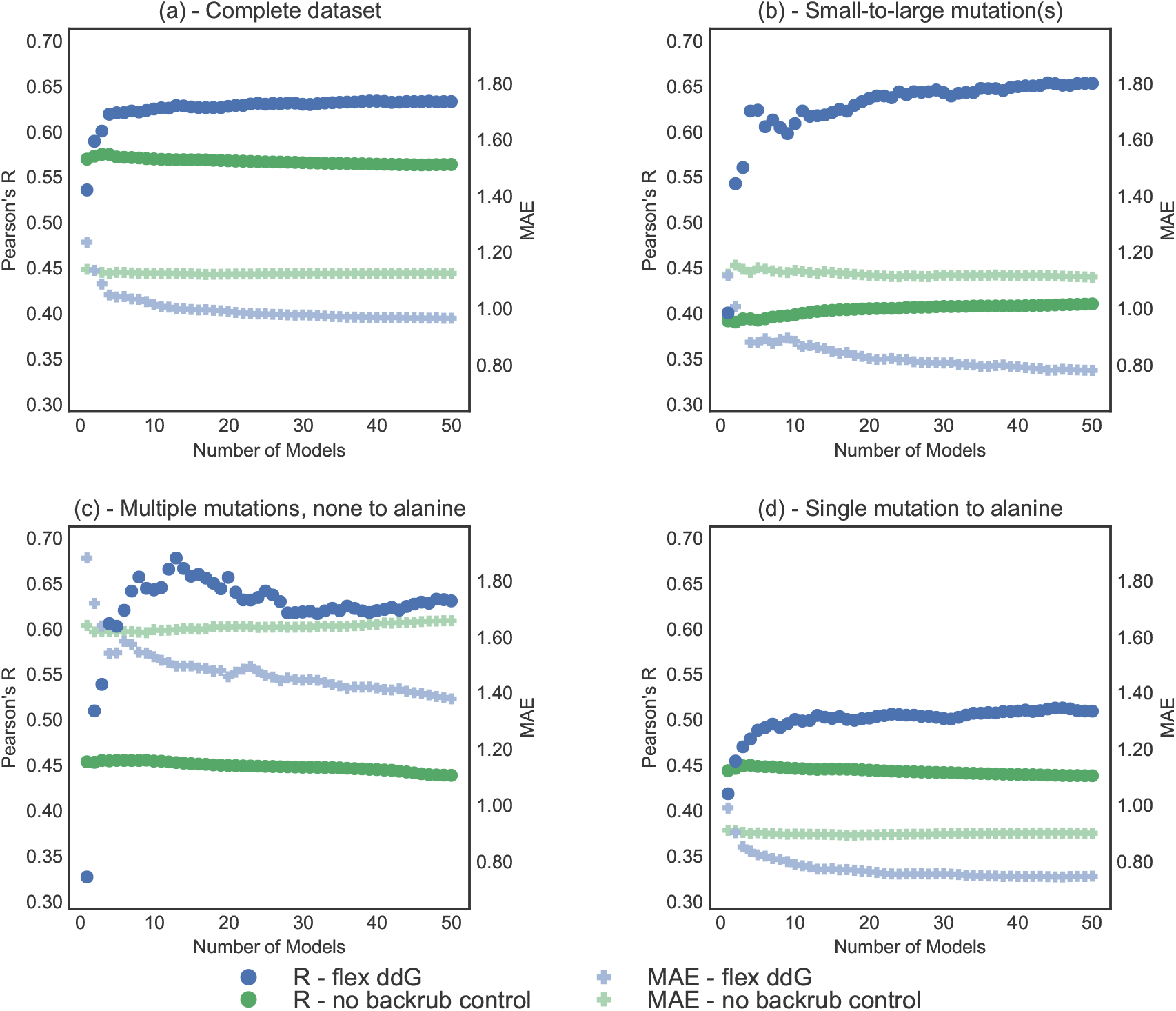
Correlation (Pearson’s R, left y-axis) and MAE (Mean Absolute Error, right y-axis) vs. number of averaged models (x-axis), on the complete ZEMu set, and subsets. Pearson’s R is shown as circles, and MAE as faded plusses. Predictions generated with the Flex ddG protocol are shown in blue. Predictions generated with the no backrub control protocol are shown in green. A selection of key data underlying this figure can be found in Table S4. Flex ddG is run with 35000 backrub steps. Structures are sorted by their minimized wild-type complex energy. (a) Complete dataset (n = 1240) (b) Small-to-large mutation(s) (n = 130) (c) Multiple mutations, none to alanine (n = 45) (d) Single mutation to alanine (n = 748).

Instead of using just the three lowest energy models,^23^ we find that the performance of the ddg_monomer method also improves as more output models are averaged (Fig. S2, Table S5). This was somewhat unexpected, as the no-backrub control method, which did not show an improvement with increasing ensemble size, is conceptually similar to the ddg_monomer method. However, the difference may arise from the fact that the ddg_monomer method ramps the weight of the repulsive Lennard-Jones term in the energy function during minimization. This strategy explores conformational space more broadly in different backbone ensemble members than minimization with a fully weighted repulsive term in the no-backrub control method. In this fashion, including more ensemble members generated by the ddg_monomer method increases the conformational plasticity sampled which in turn increases performance, as seen for the flex ddg method.

Using flex ddG, the subset of small-to-large mutations shows the largest increase in correlation with experimental ΔΔ*G* values as more models are averaged (Fig. 3(b)). This result is consistent with our reasoning above that improved modeling of conformational plasticity is important for prediction performance, and that this effect is most important for significant changes in amino acid residue size. For the subset of multiple mutations where none are mutations to alanine (Fig. 3(c)), performance overall increases substantially initially when more models are added.

Averaging across increased numbers of models also improves correlation for the subset of single mutations to alanine (Fig. 3(d)). Here, improvements are seen up to averaging about 10 models, after which performance stays approximately constant. This observation indicates that increased sampling, in the very least, is not harmful for cases where one would expect structural changes to be relatively small on average.

From a practical standpoint, generating 20-30 models should constitute sufficient sampling for most cases. Sorting the generated models by score and selecting the best scoring 20-30 out of 50 models does not appear to be necessary, as not sorting the models by score (Fig. S3, Table S6) gives similar results to sorting the models (Fig. 3).

### Effect of extent of backrub sampling in each trajectory

The extent of sampling can also be controlled by changing the number of Monte Carlo steps in the backrub simulations. Fig. 4 shows the effect of increasing the number of backrub Monte Carlo steps (while averaging all 50 models at each output step) on flex ddG performance, compared to a control method with zero backrub steps that uses only minimization and side chain packing. ΔΔ*G* scores are calculated every 2,500 backrub steps.

**Figure 4:**
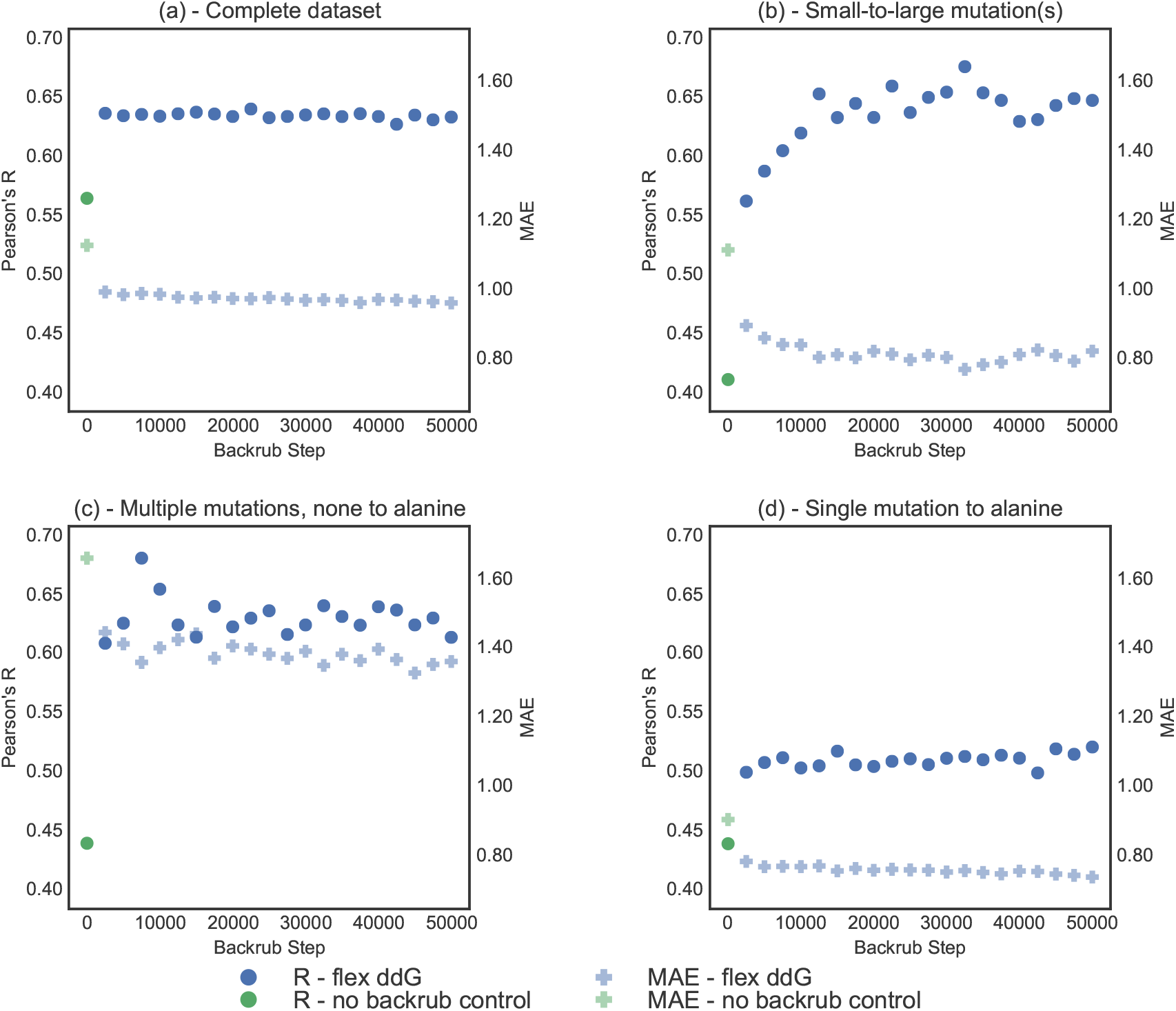
Correlation (Pearson’s R) and MAE (Mean Absolute Error) vs. number of backrub steps, on the complete ZEMu set, and subsets. Pearson’s R is shown as circles, and MAE as faded plusses. Predictions generated with the Flex ddG protocol are shown in blue. Predictions generated with the no backrub control protocol are shown in green. A selection of key data underlying this figure can be found in Table S7. (a) Complete dataset (n=1240) (b) Small-to-large mutation(s) (n=130) (c) Multiple mutations, none to alanine (n=45) (d) Single mutation to alanine (n=748)

After an initial increase for the first set of 2500 backrub steps, performance stays relatively constant for the complete dataset (Figure 4a) and for single mutations to alanine (Fig. 4d). However, for the subsets of small-to-large mutations (Figure 4b) and multiple mutations, none to alanine (Fig. 4c), performance increases considerably with increasing numbers of Monte Carlo steps. This increase in performance is similar to what was observed with averaging over more models for these subsets (Fig. 3b,c). Performance levels off at around 30,000 backrub Monte Carlo steps.

The increased performance does not appear to be simply a result of decreasing scores as the simulation progresses, as the average score of the minimized wild type complexes does not decrease uniformly across the sampled ensemble as the simulation progresses (Fig. S1). The pairwise backrub ensemble RMSDs continue to increase throughout the backrub simulation for all subsets (Fig. S4), indicating that diminishing returns at > 30,000 Monte Carlo steps is not a result of failure to sample new conformations, but rather might indicate that continued sampling does not capture additional relevant local changes in structure in this benchmark set.

### Score analysis

As the sampling and scoring problems of protein modeling are generally linked, it is often the case that improving one enables further improvements in the other.

First, we compared the performance of our flex ddG method, which was run using Rosetta’s Talaris^45,47,48^ energy function, to an identical protocol run with the more recently developed Rosetta Energy Function (REF).^52^ We did not observe an increase in performance on the complete ZEMu dataset, and performance decreases were seen for the subsets of small-to-large mutations and multiple mutations (Table S8). Interestingly, flex ddG performance with the REF energy function increased over using the Talaris energy function if the resolution of the input crystal structure was <= 1.5 Ă, but this subset of the data was rather small with only 52 mutations.

Next, we sought to analyze underlying errors of the Rosetta energy function (when applied to interface ΔΔ*G* by assessing the individual terms of the energy function. To do so, we chose to reweight the terms of the energy function using a non-linear reweighting scheme similar to Generalized Additive Models (GAMs).^51^ In this reweighting method, we used Monte Carlo sampling to fit a sigmoid function to the individual distributions of energy function terms, with the objective function of reducing the absolute error between our predictions and known experimental values over the entire dataset.

The effect on the predictions is shown in Fig. 5, Fig. S5, and Table S9. In general, the GAM-adjusted predictions contain fewer outliers. In particular, experimental ΔΔ*G* values that are relatively neutral (near zero) can sometimes be predicted by flex ddG to be highly destabilizing; the GAM model reduces the magnitude of error of many of these outliers, improving overall performance (Fig. 5). The overall correlation increases from 0.64 to 0.68 (Table 2 and Table S9) when refitting the values from the Rosetta Talaris energy function;^45,47,48^ refitting values from the Rosetta REF energy function^52^ leads to a similar increase from 0.63 to 0.68 (Fig. S5, Table S8, Table S9). The correlation coefficient also increases when refitting the values obtained for the no backrub control, but only to 0.62 (Fig. 5b, Table S9).

**Figure 5:**
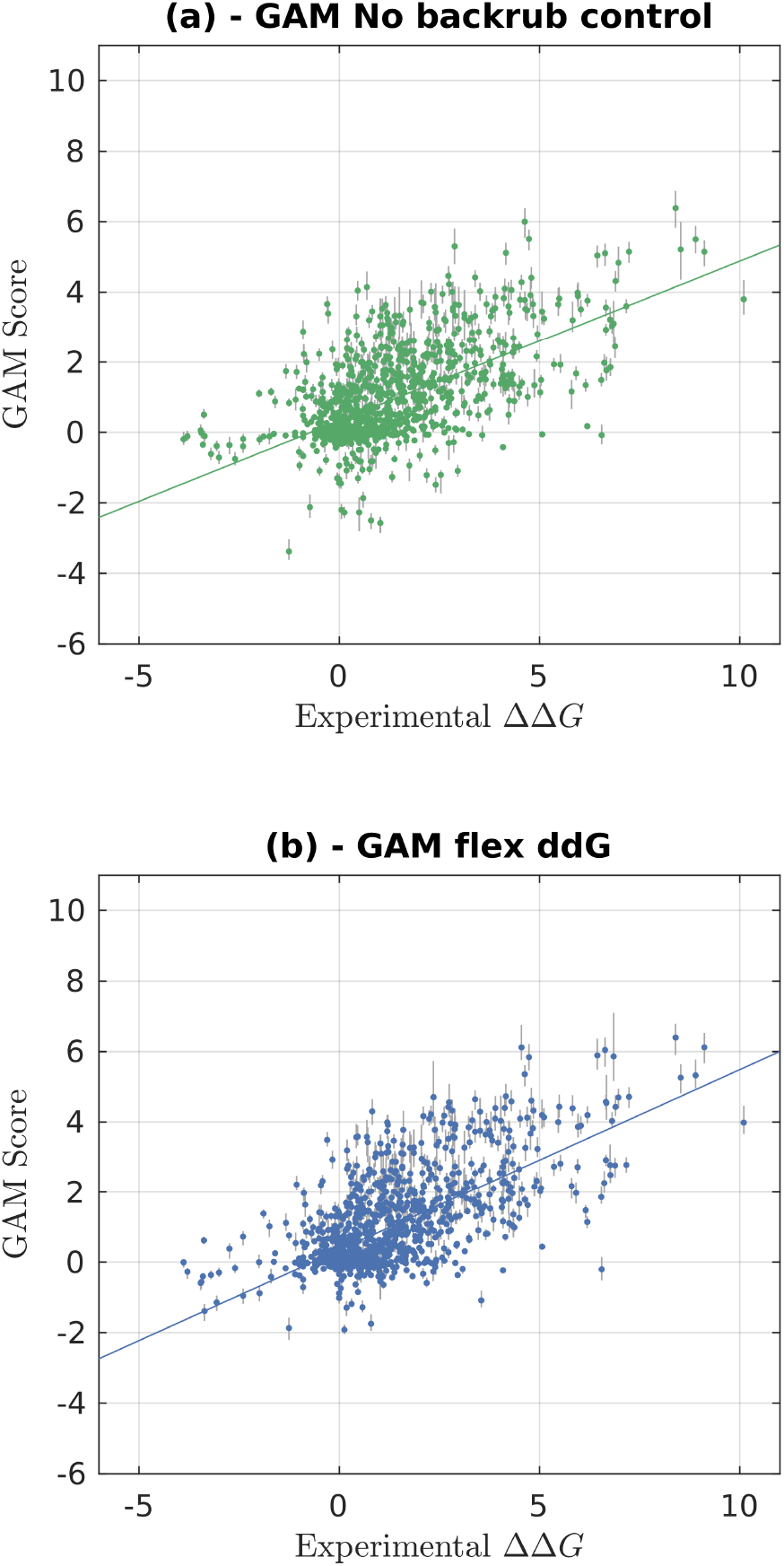
Experimentally determined ΔΔ*G* values (x-axis) versus predictions using a Generalized additive model (GAM). GAM scores are refit from values in Rosetta Energy Units (REU) using the Rosetta Talaris^45,47,48^ energy function. The error bars in gray represent the range from minimum to maximum fit predicted ΔΔ*G* value for the 1000 sampled GAM models. **(a)**: Control (no backrub) Rosetta predictions. **(b)**: Flex ddG Rosetta predictions using 35,000 backrub steps and 50 output models. A line of best fit is shown in each of the panels.

The fit functions (fit for Talaris-derived ΔΔ*G* predictions) are shown in Fig. S6. Extreme values for most score terms are downweighted, especially for the fa_sol and fa_atr terms, which make the largest contributions to predicted ΔΔ*G* (Fig. S7).

## Conclusions

We have shown on a large, curated benchmark dataset that the “flex ddG” method presented here is more accurate than previous methods for estimating changes in binding affinity after mutation in protein-protein interfaces. Particular improvement in performance is seen on the subset of small-to-large mutations, indicating that representing backbone flexibility using backrub motions is effective in cases where backbone rearrangements are expected to be more common. Other notable improvements over previous methods are seen for stabilizing mutations, mutations in antibody-antigen interfaces, and for cases with multiple changes where none of the mutations is to an alanine residue.

We have also shown that more accurate predictions can be obtained by averaging the predictions across a generated structural ensemble of backrub models, and that the number of required models is relatively low (20-30). Prior methods that produced ΔΔ*G* predictions by averaging an ensemble of models required on the order of thousands of models,^28^ indicating that backrub sampling can efficiently sample the local conformational space around an input wild-type structure that is relevant for interface ΔΔ*G* prediction.

By creating a method that uses backrub to sample conformational space more broadly than minimization alone, while still staying close to the known wild-type input structure, we have also generated data that should prove useful for future energy function improvements. In particular, using Rosetta’s newest REF energy function^52^ does not improve performance of our method when compared to use of the prior Talaris^45,47,53^ energy function (Table S8), indicating that the backrub sampling parameters might require further benchmarking and adaption to the REF energy function. Our error analysis via GAM-like reweighting also indicates potential avenues for energy function improvement by identifying imbalances in predicted energetic contributions leading to overestimation of stabilizing and destabilizing effects. Further improvements might also be obtained by more explicitly including the effects of altering water-mediated interactions^54^ and of conformational entropy,^2,55^ as well as by considering the commonly observed shortcomings of energy functions balancing the magnitudes of electrostatic interactions and desolvation costs. We expect energy function improvements to require more accurate representation of subtle conformational changes, as these changes can have a considerable impact on design predictions.^56^

## Acknowledgement

The authors acknowledge the following sources of funding: T.K. was supported by grants from the National Institute of Health (R01 GM110089 and R01 GM117189). T.K. is a Chan Zuckerberg Biohub investigator. M.H. was supported by Academy of Finland grant 299915. S.T. and J.E.L. were supported by National Science Foundation Graduate Research Fellowships.

## Supporting Information Available

The following files are available as Supporting Information:

- suppinfo.pdf: Additional figures/tables
- flex-ddG-data.csv: Rosetta scores for all predictions

---

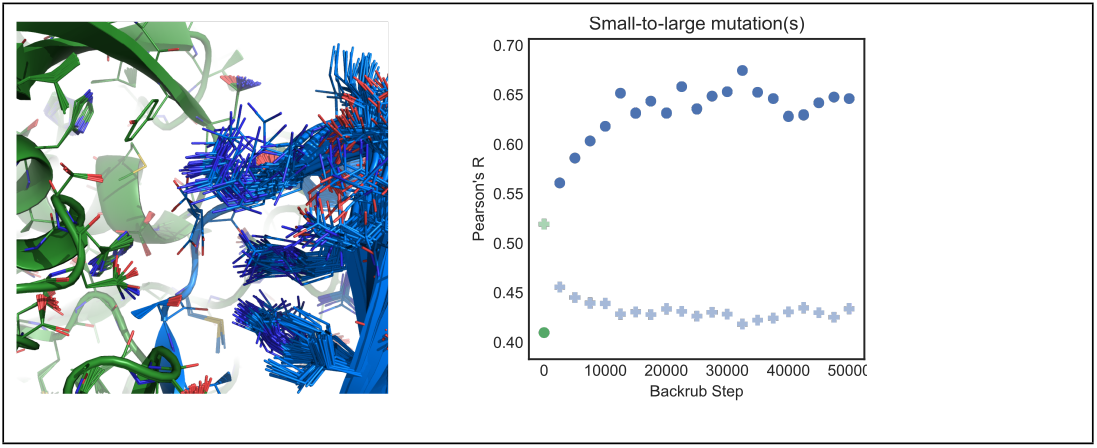
Graphical TOC Entry

